# Perceptual learning of lesions in mammograms induced by response feedback during training

**DOI:** 10.1101/752246

**Authors:** Sebastian M. Frank, Andrea Qi, Daniela Ravasio, Yuka Sasaki, Eric Rosen, Takeo Watanabe

## Abstract

Detecting subtle lesions in mammograms indicative of early breast cancer usually requires years of experience. Well-designed training paradigms could be a strong tool for promoting perceptual learning (PL) with rapid and long-lasting improvement in detectability of these subtle mammographic lesions. Given that PL occurs without feedback about the accuracy of subjects’ responses, the role of feedback has been completely ignored in clinical applications of PL. However, in this study, we found that the contents of the feedback profoundly and differentially influence the formation and retention of PL to detect calcification and architectural distortion lesions, two types of mammographic lesions that are frequently missed in mammographic screenings. We trained subjects to detect one type of lesion in a mammogram and manipulated the content of the response feedback during training for 3 groups (no feedback, correctness only, and both correctness and location of the lesion). We found that PL occurred for both lesions when both correctness and location feedback were provided. PL also occurred for calcifications but not for distortions when only correctness was provided. No learning occurred without feedback for either lesion. A retest conducted six months later showed that PL was retained only in the group with both correctness and location feedback for both types of lesions. In contrast to the general consensus of basic PL studies, our results demonstrate that the content of the response feedback is a determining factor in forming and retaining PL to detect mammographic lesions.

## Introduction

Breast cancer is the most common cancer in women and the second leading cause of cancer mortality among women in the US^1^. In fact, each year, over 250,000 cases of breast cancer are diagnosed, and 40,000 women die from breast cancer in the US. Screening mammography significantly reduces breast cancer mortality by detecting early, curable breast cancer^2^, and since its widespread utilization in the 1990s, it has resulted in approximately 250,000 fewer breast cancer deaths in the US.

However, detection of lesions indicative of early breast cancer by mammography remains challenging^3-8^. Among multiple types of mammographic lesions, it is relatively easy to detect “masses”. However, subtle manifestations of breast cancer, such as fine grouped “calcification” or “architectural distortion” lesions are known to be particularly difficult to detect without years of experience, and account for a relatively high proportion of cancers missed at screening^3, 6, 9-10^ (Figure 1). Calcifications appear as white specks tightly grouped together, while architectural distortions appear as lines radiating to a central point. Despite the importance of finding these lesions, the subtle nature of these lesions has prevented the development of a method to train young inexperienced radiologists to better detect these lesions.

**Figure 1.**
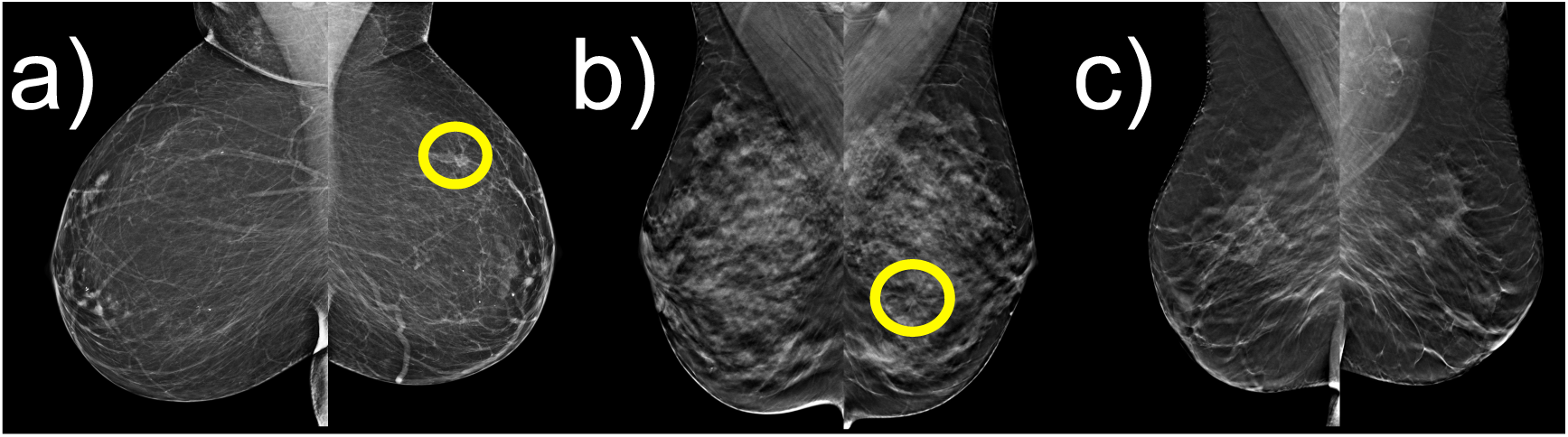
Example mammograms used for PL. Left and right breasts are presented side-by-side. The yellow circle shows the location of the lesion and was only visible during feedback (see Figure 3). (a) Calcification lesion, defined as fine white specks, tightly grouped together. (b) Architectural distortion lesion, defined as lines radiating to a central point (similar to the spokes of a wheel). (c) No lesion.

It is well known that repetitive practice of a visual task to detect a subtle difference in visual displays enhances task performance. This so-called perceptual learning (PL)^11-12^ has been studied extensively as a useful tool to improve vision. Thus, PL might also be a strong training tool to improve the detection of these subtle mammographic findings.

One distinguished aspect of PL is that it does not require feedback about the correctness of the response to a given task^13-15^. PL developed with fake feedback is rapidly modified by a few trials with correct feedback^16^. Another study shows that PL is facilitated only by positive fake feedback, which indicates that the response is correct to an incorrect response but not by negative fake feedback, which indicates that the response is incorrect to a correct response^17^. These findings led researchers to suggest that feedback merely plays a role in temporarily shifting decision criteria or enhancing the motivation of training^18-19, 17^ without changing a fundamental mechanism of PL. This has greatly discouraged researchers from examining how feedback could improve PL in clinical applications.

In contrast to such a general belief, we report here that feedback plays a fundamental role in inducing learning to better detect lesions in screening mammograms. First, we find that different contents of feedback are necessary for the PL of calcification and distortion lesions. Second, with no feedback, no PL for either lesion was observed. Third, the retention of PL for either lesion depended on the content of the feedback provided during training. These results indicate not only that feedback plays a crucial role in the formation and long-term retention of PL for mammographic lesions but also that the content of the feedback works differently on the PL of different types of lesions.

## Results

The general design of the study is shown in Figure 2. Two factors were manipulated. The first factor was the trained type of lesion (calcification lesion and architectural distortion lesion). The second factor was the type of response feedback provided during training. There were three different types of response feedback (Figure 3). In the <u>detailed feedback</u> condition, feedback about both the correctness of detection and the correctness of identification of the location of the lesion were provided. In the <u>partial feedback</u> condition, feedback only about the correctness of detection was provided. Finally, in the <u>no feedback</u> condition, no feedback was provided. Each subject was assigned to a combination of a trained lesion and type of response feedback during training. There was a total of n = 12 subjects for each combination of trained lesion and response feedback (for a total of n = 72 subjects). Subjects were college-age students with no medical background and without any prior exposure to mammographic images (see Methods for details).

**Figure 2.**
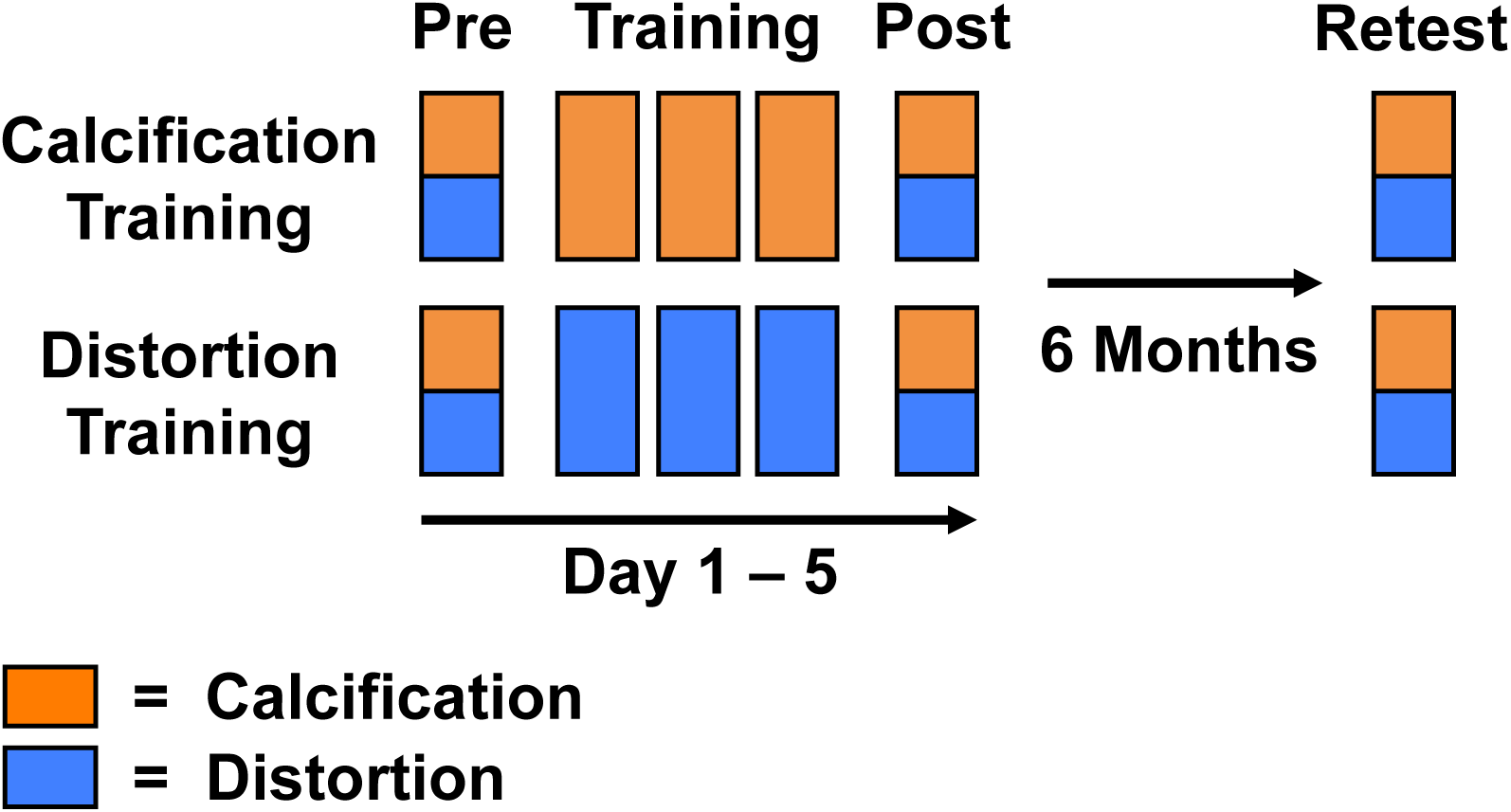
Training procedures. A total of six sessions were conducted on separate days. During the pretest, posttest and retest, subjects were tested on both types of lesions without any response feedback. Between the pretest and posttest, there were three training sessions during which different groups of subjects were trained either on calcifications (orange) or architectural distortions (blue) with detailed feedback, partial feedback, or no feedback (see Figure 3). The retest was conducted on a subset of subjects six months after the end of original training.

**Figure 3.**
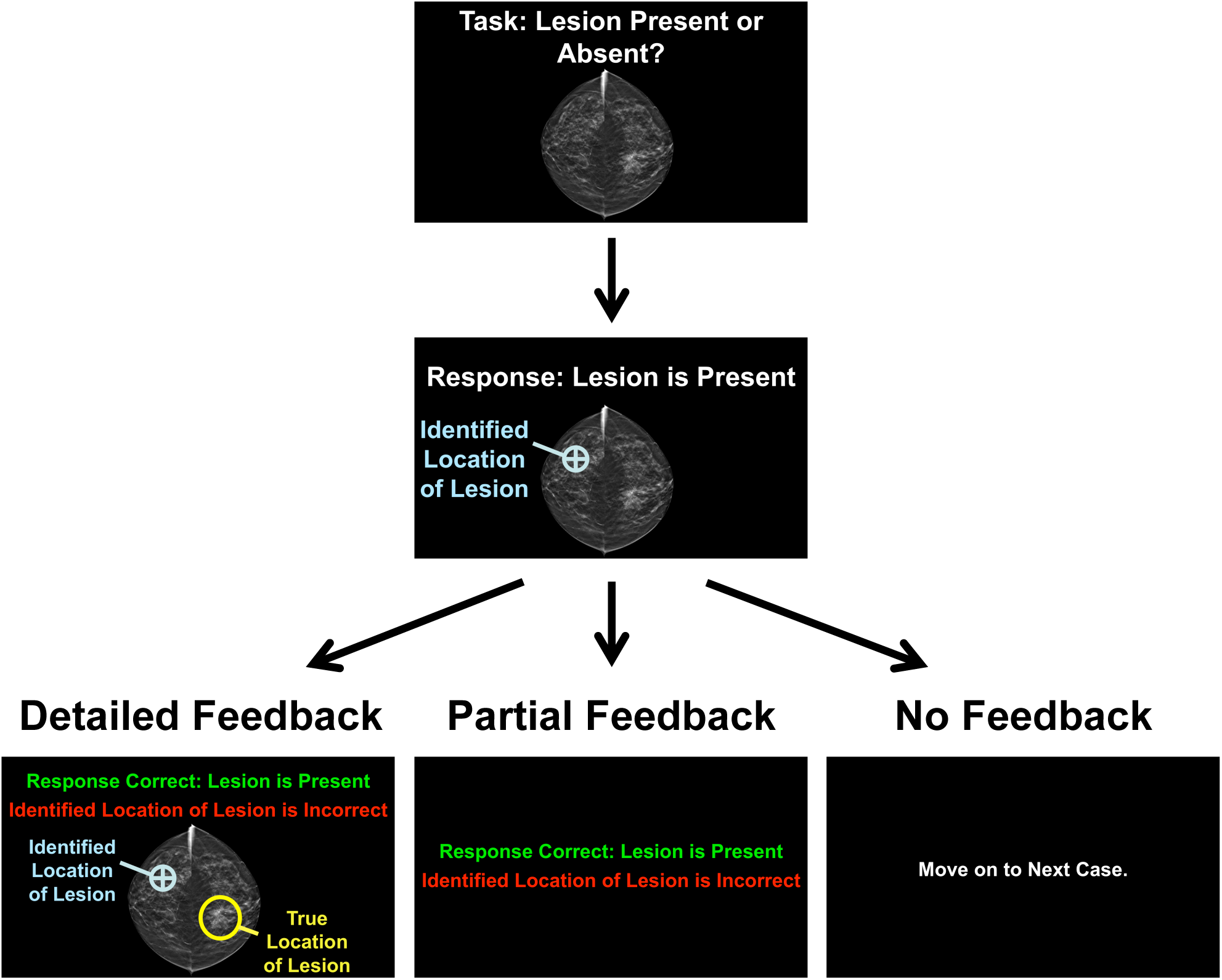
Examples of training trials. Subjects were presented with a mammogram and asked to indicate the presence or absence of a lesion by pressing one of two buttons on the keyboard. There was no time limit for their response. If subjects reported the presence of a lesion, they were further asked to indicate the center of the lesion by moving and clicking the cursor. The selected location was then highlighted by a blue crosshair. Depending on the assigned feedback condition, subjects were then presented with different response feedback conditions. In the detailed feedback condition (bottom left), subjects were informed about the correctness of their response and the true location of the lesion. The mammogram was then shown again, and the true location of the lesion was enclosed by a yellow circle. Subjects could compare the true location with the location they had identified (denoted by blue crosshair). If there was no lesion, the mammogram was presented without any yellow circle. In the partial feedback condition (bottom center), subjects were only informed about the correctness of their response. In the no feedback condition (bottom right), subjects did not receive any response feedback and moved on to the next trial.

Each subject performed pre- and posttests on both lesions. Thus, both trained and untrained lesions were tested before and after training. These tests were conducted without any feedback. On each of the pre- and posttest trials, subjects were asked to indicate whether a lesion was present or absent in the mammogram by pressing one of two buttons on a keyboard. If they reported the presence of a lesion, subjects were further asked to locate the lesion in the mammogram by moving the cursor to the center of the lesion and clicking on the center. Finally, subjects identified which type of lesion it was (calcification or distortion) by pressing one of two buttons on the keyboard. Mammograms with lesions were presented intermixed with mammograms without any lesion. If a lesion was present, the subject’s response was considered correct (hit) if the subject correctly detected, located, and identified the type of lesion. If a lesion was absent, the subject’s response was considered correct if the subject correctly rejected the presence of a lesion (correct rejection). Performance was computed as observer sensitivity (d’). A total of 50 trials were conducted in each test session (20% of the trials contained a mammogram with a calcification lesion, another 20% contained a mammogram with a distortion lesion, and the remaining 60% were mammograms without any lesion; see Methods for details).

In between the pretest and posttest, three training sessions were conducted on separate days. In each of these sessions, subjects were trained on their assigned combination of lesion and response feedback. For each trial, subjects were asked to indicate whether a lesion was present and, if so, where it was located, in a similar manner to the pretest and posttest. During training, only mammograms with the trained lesion type were presented intermixed with mammograms without any lesions. Subjects were informed about their assigned lesion for training and thus were not asked to indicate the type of lesion as part of their response during training. If a lesion was present, the subject’s response was considered correct (hit) if the lesion was correctly detected and located. If a lesion was absent, the subject’s response was considered correct if the subject rejected the presence of a lesion. A total of 50 trials were conducted in each training session (in 20% of the trials, the mammogram contained a lesion).

A subset of subjects (n = 43 in total) was available for a retest that was conducted six months after the end of original training, with exactly the same procedures as those of the pretest and posttest. See Methods for more details.

### Perceptual Learning of Trained Lesion

Figure 4 shows the mean performance (corresponding to observer sensitivity, d’) across subjects for the pretest and posttest (see Supplementary Figure 1 for performance in training sessions and Supplementary Figure 2 for hit and false-alarm rates). A 2 × 2 x 3 mixed ANOVA with a within-subject factor of session (pre, post) and between-subject factors of trained type of lesion (calcification, distortion) and content of response feedback (detailed, partial, none) showed a significant three-way interaction of session, trained type of lesion and feedback [*F*(2,66) = 3.16, *p* = 0.049, partial *η*^2^ = 0.09], suggesting that the content of response feedback influenced PL differently for different lesions. Furthermore, there was a significant two-way interaction between session and feedback [*F*(2,66) = 6.21, *p* = 0.003, partial *η*^2^ = 0.16]. The ANOVA also yielded a significant main effect of trained type of lesion [*F*(1,66) = 58.0, *p* < 0.001, partial *η*^2^ = 0.47], suggesting that d’ was greater for calcifications than distortions across sessions and feedback conditions (Figure 4). There was a significant main effect of session [*F*(1,66) = 30.4, *p* < 0.001, partial *η*^2^ = 0.32], indicating that d’ was greater after training across different feedback conditions. No other main effect or interaction was significant.

**Figure 4.**
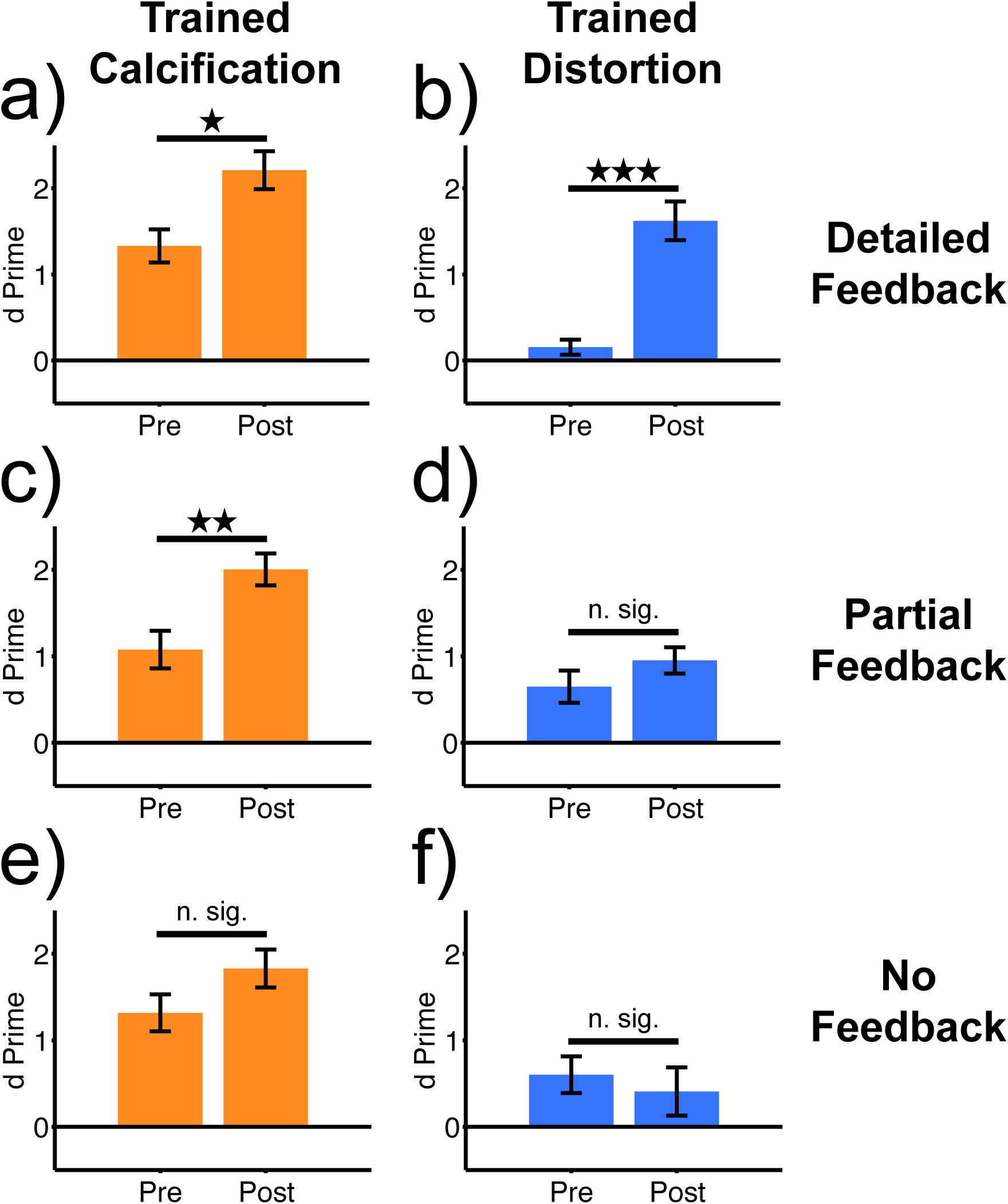
Results of the experiment. Pre and Post on the x-axis correspond to the pretest and posttest, which were conducted before and after training, respectively. Orange bars represent the mean (± SE) observer sensitivity (d’) for calcifications in subjects trained on calcifications (n = 12 for each feedback condition). Blue bars represent the mean (± SE) observer sensitivity for distortions in subjects trained on distortions (n = 12 for each feedback condition). (a) and (b): with detailed feedback during training. (c) and (d): with partial feedback during training. (e) and (f): with no feedback during training. Asterisks show the results of post hoc paired sample *t*-tests between pre- and posttests (*** *p* < 0.001, ** *p* < 0.01, * *p* < 0.05, n. sig. = no significant difference).

Post hoc *t*-tests were conducted to illustrate the relationship among trained lesion, content of response feedback and PL from pre- to posttesting. In the detailed feedback condition, PL occurred from pre- to posttesting for both calcifications [*t*(11) = 2.80, *p* = 0.02, *d* = 1.23] and distortions [*t*(11) = 5.98, *p* < 0.001, *d* = 2.49]. However, in the partial feedback condition, PL occurred only for calcifications [*t*(11) = 3.18, *p* = 0.009, *d* = 1.33] but not for distortions [*t*(11) = 1.09, *p* = 0.30, *d* = 0.51]. In the training condition without feedback, there was no PL for either calcifications [*t*(11) = 1.64, *p* = 0.13, *d* = 0.68] or distortions [*t*(11) = −0.68, *p* = 0.51, *d* = −0.23].

### Retention of Perceptual Learning for Trained Lesions

We conducted a retest six months after the end of original training with the subset of subjects from training conditions with PL who were available for retesting (Figure 5; see Supplementary Figure 3 for hit and false alarm rates). If subjects completely retained PL, we expected that d’ would not be significantly different between retest and posttest but would be significantly greater during the retest compared with the pretest. The results showed that in the detailed feedback condition, the d’ for calcifications in the retest was significantly greater than in the pretest [*t*(8) = 2.41, *p* = 0.04, *d* = 1.35] but not significantly different from the posttest [*t*(8) = −1.02, *p* = 0.34, *d* = −0.10] (Figure 5a). For distortions, although the d’ in the retest was significantly greater than in the pretest [*t*(8) = 3.52, *p* = 0.008, *d* = 1.57], it was significantly smaller than in the posttest [*t*(8) = −4.47, *p* = 0.002, *d* = −0.70] (Figure 5b). These results suggest that in the detailed feedback conditions, PL for calcifications was completely retained after six months, while PL for distortions was partially retained.

**Figure 5.**
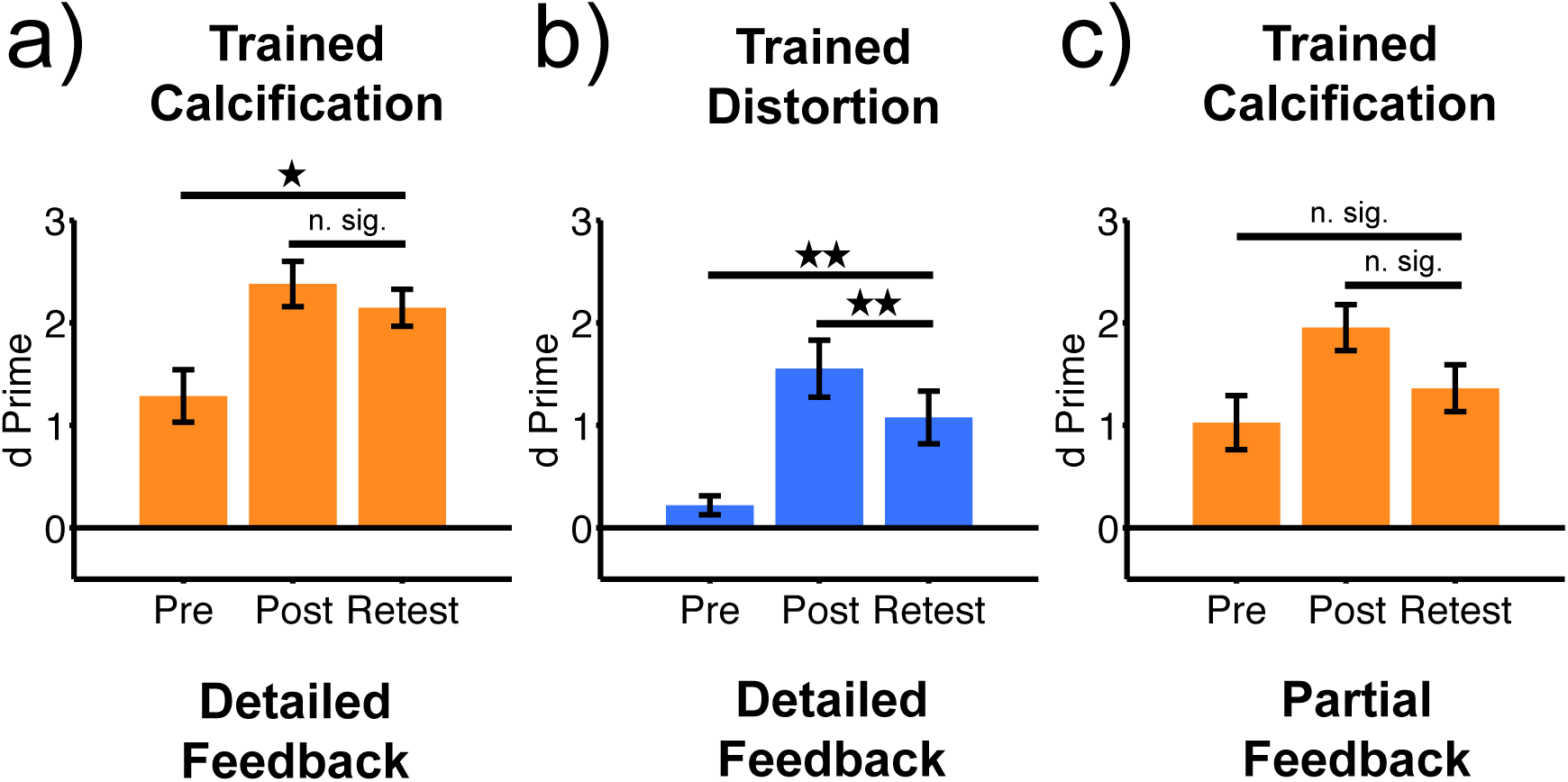
Retention of PL. Pre and Post on the x-axis represent the pretest and posttest, respectively. The retest was conducted six months after the posttest. Orange bars represent the mean (± SE) observer sensitivity (d’) for calcifications in subjects who were trained on calcifications and participated in the retest (n = 9 subjects in the detailed and n = 9 subjects in the partial feedback training conditions). Blue bars represent the mean (± SE) observer sensitivity for distortions in subjects who were trained on distortions and participated in the retest (n = 9 subjects in the detailed feedback condition). (a) and (b): with detailed feedback during training. (c): with partial feedback during training. Asterisks show the results of post hoc paired sample *t*-tests between retest and pre/posttests (** *p* < 0.01, * *p* < 0.05, n. sig. = no significant difference).

PL also occurred for calcifications with partial feedback (Figure 4). However, the d’ for calcifications in the retest was not significantly different from the d’ in either the pretest [*t*(8) = 1.17, *p* = 0.28, *d* = 0.40] or the posttest [*t*(8) = −1.45, *p* = 0.19, *d* = −0.97] (Figure 5c). This suggests that PL for calcifications in the partial feedback training condition was not retained after six months.

### Perceptual Learning of Untrained Lesions

We also analyzed whether observer sensitivity changed for the untrained lesion in each training group (that is, performance on distortions in the pre- and posttests for subjects who trained on calcifications and vice versa, see Supplementary Figures 4 and 5). A 2 × 2 x 3 mixed ANOVA with the within-subject factor of session (pre, post) and between-subject factors of untrained type of lesion (calcification, distortion) and content of response feedback for the trained lesion (detailed, partial, no feedback) was conducted. Importantly, there was no significant interaction between session, untrained type of lesion and content of the response feedback [*F*(2,66) = 0.61, *p* = 0.55, partial *η*^2^ = 0.02]. Furthermore, there were no significant interactions between session and content of the response feedback [*F*(2,66) = 0.19, *p* = 0.83, partial *η*^2^ = 0.01] or between untrained type of lesion and content of the response feedback [*F*(2,66) = 0.21, *p* = 0.81, partial *η*^2^ = 0.01]. There was no significant main effect of content of the response feedback [*F*(2,66) = 0.73, *p* = 0.49, partial *η*^2^ = 0.02]. Taken together, these results suggest that the content of the response feedback given for the trained type of lesion during training did not influence d’ in any session for any type of untrained lesions.

The ANOVA yielded a significant interaction between session and untrained type of lesion [*F*(1,66) = 5.93, *p* = 0.02, partial *η*^2^ = 0.08], suggesting that d’ was greater for untrained calcifications compared with untrained distortions in the posttest compared with the pretest (Supplementary Figure 4). Furthermore, there was a significant main effect of session [*F*(1,66) = 5.97, *p* = 0.02, partial *η*^2^ = 0.08], suggesting that d’ for untrained lesions was better in the posttest compared with the pretest. Moreover, there was a significant main effect of untrained type of lesion [*F*(1,66) = 25.8, *p* < 0.001, partial *η*^2^ = 0.28], indicating that d’ on untrained calcifications was superior to untrained distortions across tests (Supplementary Figure 4).

### Retention of Perceptual Learning for Untrained Lesion

The retention of improvements in d’ for untrained calcifications was examined in the retest (Supplementary Figures 6 and 7). Since the comparison between pretest and posttest did not indicate any influence of the response feedback on the untrained lesions (see above), data from the subjects were combined across feedback conditions. The results show that the d’ for untrained calcifications during retest was not significantly different from that from either pretest [*t*(24) = 1.12, *p* = 0.27, *d* = 0.21] or posttest [*t*(24) = −1.15, *p* = 0.26, *d* = −0.39], indicating that PL for untrained calcifications from pre- to posttest (see above) was not retained after six months.

## Discussion

In this study, we investigated how response feedback during training influences the PL of lesions in mammograms. Naïve subjects were trained on either calcification or architectural distortion lesions, both of which are indicative of early breast cancer and are commonly missed in mammogram screenings^3-4, 7^. During training, the content of the response feedback was varied between subjects. In training with detailed feedback, subjects were informed about the correctness of their response and about the true location of the lesion. In training with partial feedback, subjects only learned about the correctness of their response but could not review the true location of the lesion. A third group of subjects trained without any response feedback. The results of our study suggest that the content of the response feedback influenced the PL of calcifications and distortions differently. The PL of calcifications was already observed in training with partial feedback, whereas the PL of distortions only occurred in training with detailed feedback. However, the PL for both lesions showed long-term retention only in training with detailed feedback.

PL of low-level visual features can occur without external feedback during training^13-15^. In addition, the effects of feedback on PL are short-lasting^16^. These results have led researchers to generally believe that feedback does not play a fundamental role in PL. To our knowledge, no research has been conducted to systematically determine the role of feedback in PL in clinical applications. In contrast to the general belief that feedback is of little importance to PL, the results of the present study indicate that feedback is indispensable to the PL of lesions in mammograms and that the choice of the right type of feedback makes visual training highly effective and long-lasting for improving the detectability of the trained lesion.

These results include the following findings: (1) no PL was observed for either calcification or distortion lesions when subjects trained without feedback; (2) PL for calcifications could occur when subjects were only informed about the correctness of their response after each trial without any opportunity to review the examined mammogram one more time (partial feedback); (3) in contrast, significant improvements in the detection of distortions were only observed when subjects reviewed the mammogram and the true location of the lesion (detailed feedback); and (4) long-lasting improvements in detecting calcification and distortion lesions were only induced by detailed feedback during training. Taken together, these results indicate that response feedback plays a fundamental role in inducing a long-lasting PL of lesions in mammographic screenings.

In basic and clinical studies in which feedback is used to visually train subjects, partial feedback^20-21^ or detailed feedback^22-23^ has never been used systematically. Our results suggest that training using partial feedback is sufficient for PL of calcifications. However, training using partial feedback is not sufficient for PL of distortions. This difference in the effectiveness of partial feedback might be related to differences in the difficulty of detecting calcifications and distortions. Specifically, our results show that subjects achieved better detection performance for calcifications compared with distortions before training (see Figure 4). Thus, the effectiveness of the response feedback might be associated with the difficulty of detecting different types of lesions. However, contrary to this assumption, we found that PL was not long-lasting without detailed feedback, neither for calcifications nor for distortions. This indicates that detailed feedback plays a crucial role, at least in long-term PL, for both calcifications and distortions, irrespective of their detection difficulty.

We found that the effects of feedback were specific to the PL of the trained type of lesion and did not transfer to the untrained type of lesion. This is in accord with the hypothesis that the PL of mammographic lesions occurs at a relatively low stage of visual processing. Recently, methods to eliminate the location specificity of PL have been developed^24-25^. The integration of adequate feedback with these methods may help generalize the PL of mammographic lesions to a greater degree.

Why does detailed feedback induce a stable PL? We speculate that the opportunity for subjects to review their responses together with the correct response after each training trial might serve as an “instructor” to PL. Although subjects were familiarized with the features that characterize different mammographic lesions before training, the large degree of individual variation of the trained lesion in mammograms from different patients renders the creation of a general search template for calcification or distortion lesions difficult. Feedback after each trial with information about the true location of the lesion might facilitate the creation of such a search template over the course of training and thus induce stable and long-lasting PL.

To conclude, PL with response feedback significantly improves the detectability of subtle lesions in mammograms by naïve subjects. We found that feedback about the correctness of the response improves the detectability of calcifications (partial feedback), while training with feedback about the location of a lesion as well as the correctness of the response (detailed feedback) is necessary for the development of PL of distortion lesions. However, long-lasting improvements for both lesions depend on detailed feedback. Thus, the content of the response feedback plays a crucial role in the PL of mammographic findings. The results of this study indicate the importance of future studies on the effect of the content of feedback on PL in basic research and clinical investigations.

## Methods

### Subjects

A total of n = 72 subjects [average age (± SE) = 21.1 ± 0.44 years, 44 females, 66 right-handers] participated in the study. Subjects gave informed, written consent prior to participation. The study was approved by the local ethics board at Brown University and the University of Colorado, Denver.

### Breast Images

Two-dimensional grayscale mammographic images obtained from routine screening examinations on asymptomatic women performed using Hologic Selenia units at the University of Colorado, Denver, were used for the study. Images were presented at a resolution of 2000 × 1125 pixels. Each image was evaluated by an expert radiologist (E.R., 22 years of experience) and classified as normal or as having either a calcification or distortion lesion. Images consisted of either a single 2D image or a single 1 mm thick tomosynthesis slice and were deemed representative of either the lesion or normal findings by E.R. Each mammogram was from a different patient. In the majority of cases (286 out of 330 mammograms in total), both breasts of the patient were scanned and presented side-by-side to subjects in the experiment. The remaining 14 mammograms were only collected for one breast. If there was a lesion, it was only present in one breast. The image background was black, and no additional preprocessing was performed on the mammograms. There was a pool of 60 mammograms with calcification lesions, 60 mammograms with distortion lesions, and 210 normal mammograms. The same pool of images was used for each subject. Images were randomly sorted into one of the six sessions for each subject.

### Experimental Schedule

The study consisted of six sessions, which were conducted on separate days (Figure 2). In the first session (pretest), detection performance for calcifications and distortions was measured prior to training in each subject. No feedback about performance was provided during the test. In the second, third, and fourth sessions, subjects were trained either on calcification or distortion lesions (depending on group assignment, see below) and received feedback after each training trial (depending on the feedback condition, see below). In the fifth session (posttest), performance after training was measured as in the pretest for both the trained and untrained lesions. On average (± SE), 6.34 ± 0.26 months after the posttest, the retention of PL was measured in a retest session similar to the pre- and posttests for both the trained und untrained lesions. All subjects (n = 72) participated in the training and the pre- and posttests. There was a total of n = 12 subjects for each combination of a trained lesion and feedback during training. A subset of trained subjects (n = 43) was available for the retest.

### Pretest/Posttest/Retest

A total of 50 mammograms were presented to the subjects in each test. Ten mammograms contained a calcification lesion, another ten contained a distortion lesion, and the remaining mammograms did not contain any lesions. Mammograms were presented in a random order. On each trial, subjects were asked to indicate whether a lesion was present in the mammogram by pressing one of two buttons for “yes” and “no” on a keyboard. If subjects indicated the presence of a lesion, they were further asked to locate the center of the lesion by a mouse-click. Finally, subjects were asked to report whether the lesion was a calcification or a distortion by pressing one of two buttons on the keyboard. If subjects reported that no lesion was present, they moved on to the next case.

Subjects could move each mammogram to left and right or up and down. Moreover, they could zoom into the mammogram to magnify anatomical details or zoom out. Subjects could search the mammogram overtly by making eye movements. If they located a lesion, the subjects could change their decision until making a final confirmation of their selection. The experiment was self-paced with no time limit. No feedback was provided. The completion of a test session took ∼ 30 min. Subjects were seated 60 cm away from the computer screen (size: 43° x 27°). No chin rest was used. Room lights were turned off. Stimuli were presented with the Psychophysics Toolbox^26-27^ running in MATLAB (The MathWorks, Natick, MA). Prior to each test session, instruction slides with example mammograms with distortion and calcification lesions were presented to the subjects.

### Training

There were two training groups (calcification and distortion lesions) and three different feedback conditions (detailed, partial, or no feedback). Each subject was randomly assigned to a combination of a trained lesion and feedback condition.

Training sessions were similar to test sessions except that 10 mammograms with either calcification or distortion lesions (dependent on the training group) were included along with 40 mammograms without any lesion. Subjects were informed about their assignment to either calcification or distortion lesions training prior to the first training session. On each training trial, subjects were asked to indicate whether the assigned lesion was present in the mammogram, and if so, they were further asked to report where the center of the lesion was located by moving the cursor and clicking on the center. Detailed feedback, partial feedback, or no feedback was provided at the end of each trial. In the detailed feedback condition, subjects were informed about the correctness of their response as well as the true location of the lesion. This was done by providing a written statement about the correctness of the response to the subjects in green (for a correct response) or red (for an incorrect response) (Figure 3a). Depending on the correctness of the subject response, this statement could read: “Response correct: lesion is present”. “Response correct: lesion is not present”. “Response incorrect: lesion is present”. “Response incorrect: lesion is not present”. The mammogram was then presented again, and the location of the lesion identified by the subject as well as the true location of the lesion (if it was present) were indicated. A blue crosshair indicated the location of the center of the lesion as identified by the subject, and a yellow circle enclosed the true location of the whole lesion (if a lesion was present; if no lesion was present, only the blue crosshair with the subject response was shown). Subjects could magnify or move the mammogram in different directions for review. In the partial feedback condition, only feedback about the correctness of the response was provided to the subjects. As in the detailed feedback condition, a written statement was presented at the end of the trial (Figure 3b). In the no feedback condition, subjects did not learn about the correctness of their response at the end of the trial (Figure 3c) and were only requested to press a button to move on to the next trial. Each training session was self-paced and took ∼ 30 min.

### Analysis

Performance in test and training sessions was quantified as observer sensitivity (d’). During pretest, posttest and retest, the subject’s response was considered a hit when the following three conditions were all met: (1) the subject correctly identified the presence of a lesion, (2) the center of the lesion as identified by the subject was within the area corresponding to the true location of the lesion and (3) the subject correctly identified whether the lesion was a calcification or an architectural distortion. If one of the three conditions was not met or if the subject incorrectly indicated the absence of a lesion, the response was considered a miss. If the subject incorrectly indicated the presence of a lesion, the subject response was considered a false alarm. If no lesion was present and the subject correctly indicated the absence of a lesion, the subject response was considered a correct rejection. Hit and false alarm rates were computed separately for calcifications and distortions. For training sessions, hits corresponded to trials where subjects correctly indicated the presence of the trained lesion and correctly identified its location.

### Statistics

Data were analyzed using parametric statistics (ANOVA and two-tailed *t*-tests). Measures of effect size (partial *η*^2^ for ANOVAs and Cohen’s *d* for *t*-tests) were reported.

## Acknowledgment

This work was supported by NIH R21EY028329, NIH R01EY019466, NIH R01EY027841, United States – Israel Binational Science Foundation BSF2016058

## Author Contributions

S.M.F., E.R., T.W. designed the study. E.R. provided and classified mammograms. S.M.F., A.Q., D.R. collected data. S.M.F. analyzed the data. S.M.F., E.R., Y.S., T.W. wrote the manuscript.

## References

1. SEER cancer data (2019). https://seer.cancer.gov/

2. Tabár, L., et al. The incidence of fatal breast cancer measures the increased effectiveness of therapy in women participating in mammography screening. Cancer 125, 515–523 (2019).

3. Bird, R. E., Wallace, T. W. & Yankaskas, B. C. Analysis of cancers missed at screening mammography. Radiology 184, 613–617 (1992).

4. Goergen, S. K., Evans, J., Cohen, G. P. & MacMillan, J. H. Characteristics of breast carcinomas missed by screening radiologists. Radiology 204, 131–135 (1997).

5. Huynh, P. T., Jarolimek, A. M. & Daye, S. The false-negative mammogram. Radiographics 18, 1137–1154 (1998).

6. Birdwell, R. L., Ikeda, D. M., O’Shaughnessy, K. F. & Sickles, E. A. Mammographic characteristics of 115 missed cancers later detected with screening mammography and the potential utility of computer-aided detection. Radiology 219, 192–202 (2001).

7. Majid, A. S., de Paredes, E. S., Doherty, R. D., Sharma, N. R. & Salvador, X. Missed breast carcinoma: pitfalls and pearls. Radiographics 23, 881–895 (2003).

8. Elmore, J. G., et al. Variability in interpretive performance at screening mammography and radiologists’ characteristics associated with accuracy. Radiology 253, 641–651 (2009).

9. Rangayyan, R. M., Banik, S. & Desautels, J. L. Computer-aided detection of architectural distortion in prior mammograms of interval cancer. Journal of Digital Imaging 23, 611–631 (2010).

10. Bahl, M., Baker, J. A., Kinsey, E. N. & Ghate, S. V. Architectural distortion on mammography: correlation with pathologic outcomes and predictors of malignancy. American Journal of Roentgenology 205, 1339–1345 (2015).

11. Sasaki, Y., Nanez, J. E. & Watanabe, T. Advances in visual perceptual learning and plasticity. Nature Reviews Neuroscience 11, 53–60 (2010).

12. Watanabe, T. & Sasaki, Y. Perceptual learning: toward a comprehensive theory. Annual Review of Psychology 66, 197–221 (2015).

13. Karni, A. & Sagi, D. Where practice makes perfect in texture discrimination: evidence for primary visual cortex plasticity. Proceedings of the National Academy of Sciences 88, 4966–4970 (1991).

14. Karni, A. & Sagi, D. The time course of learning a visual skill. Nature 365, 250–252 (1993).

15. Fahle, M. & Edelman, S. Long-term learning in vernier acuity: Effects of stimulus orientation, range and of feedback. Vision Research 33, 397–412 (1993).

16. Herzog, M. H. & Fahle, M. Effects of biased feedback on learning and deciding in a vernier discrimination task. Vision Research 39, 4232–4243 (1999).

17. Shibata, K., Yamagishi, N., Ishii, S. & Kawato, M. Boosting perceptual learning by fake feedback. Vision Research 49, 2574–2585 (2009).

18. Petrov, A. A., Dosher, B. A. & Lu, Z. L. Perceptual learning without feedback in non-stationary contexts: Data and model. Vision Research 46, 3177–3197 (2006).

19. Herzog, M. H. & Fahle, M. Modeling perceptual learning: Difficulties and how they can be overcome. Biological Cybernetics 78, 107–117 (1998).

20. Sowden, P. T., Davies, I. R. & Roling, P. Perceptual learning of the detection of features in X-ray images: a functional role for improvements in adults’ visual sensitivity? Journal of Experimental Psychology: Human Perception and Performance 26, 379–390 (2000).

21. Xu, B., Rourke, L., Robinson, J. K. & Tanaka, J. W. Training melanoma detection in photographs using the perceptual expertise training approach. Applied Cognitive Psychology 30, 750–756 (2016).

22. Krasne, S., Hillman, J. D., Kellman, P. J. & Drake, T. A. Applying perceptual and adaptive learning techniques for teaching introductory histopathology. Journal of Pathology Informatics 4 (2013).

23. Chen, W., HolcDorf, D., McCusker, M. W., Gaillard, F. & Howe, P. D. Perceptual training to improve hip fracture identification in conventional radiographs. PloS one 12, e0189192 (2017).

24. Harris, H., Gliksberg, M. & Sagi, D. Generalized perceptual learning in the absence of sensory adaptation. Current Biology 22, 1813–1817 (2012).

25. Xiao, L. Q., et al. Complete transfer of perceptual learning across retinal locations enabled by double training. Current Biology 18, 1922–1926 (2008).

26. Brainard, D. H. The psychophysics toolbox. Spatial Vision 10, 433–436 (1997).

27. Pelli, D. G. The VideoToolbox software for visual psychophysics: Transforming numbers into movies. Spatial Vision 10, 437–442 (1997).

