## Supplementary Material for "Perceptual learning of lesions in mammograms induced by response feedback during training"

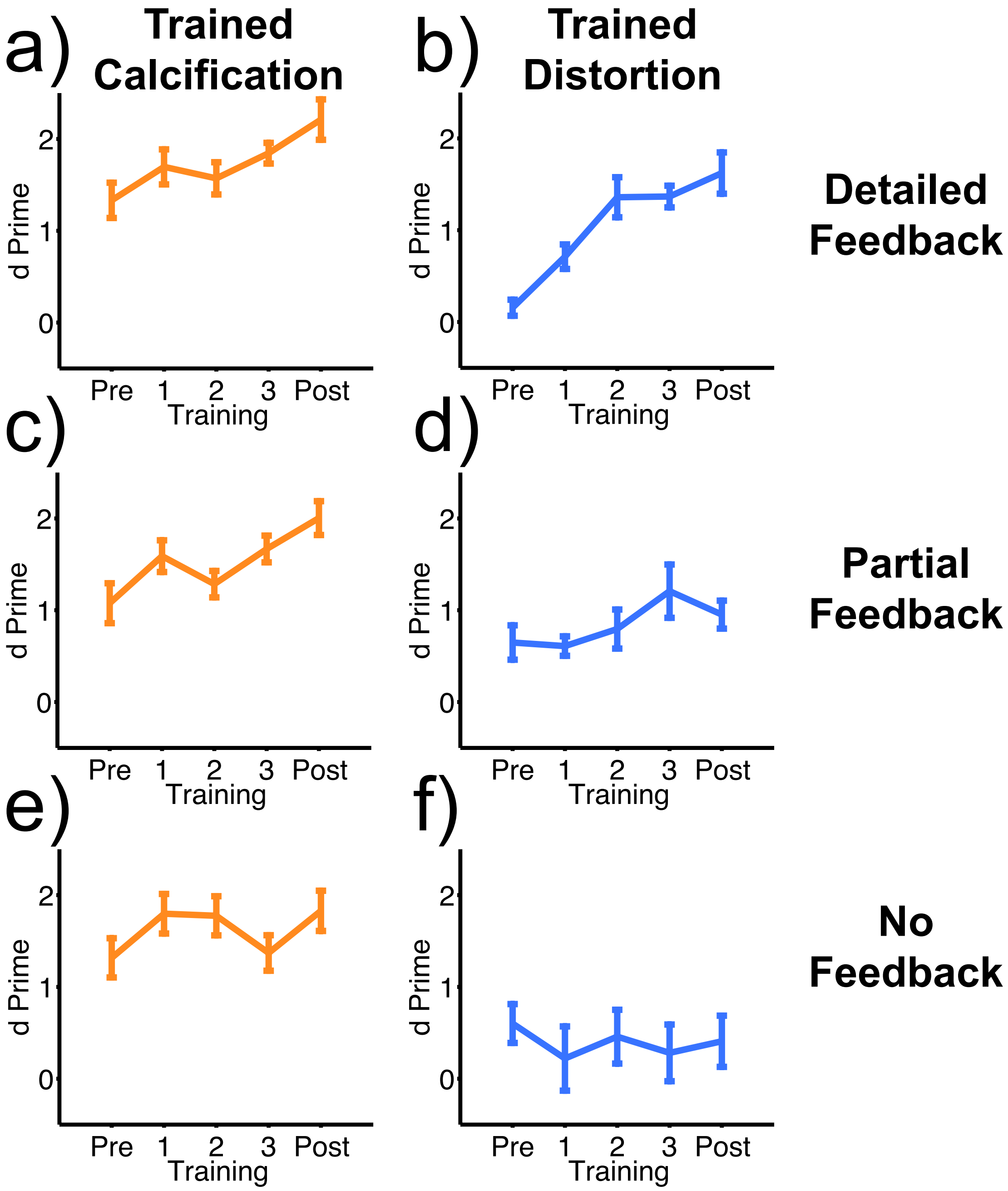


**Supplementary Figure 1**

PL for trained lesions across pretest (Pre), three training sessions and posttest (Post). Orange lines represent the mean (± SE) observer sensitivity (d’) for calcifications in subjects trained on calcifications (n = 12 for each feedback condition). Blue lines represent the mean (± SE) observer sensitivity for distortions in subjects trained on distortions (n = 12 for each feedback condition). (a) and (b): with detailed feedback during training. (c) and (d): with partial feedback during training. (e) and (f): with no feedback during training.

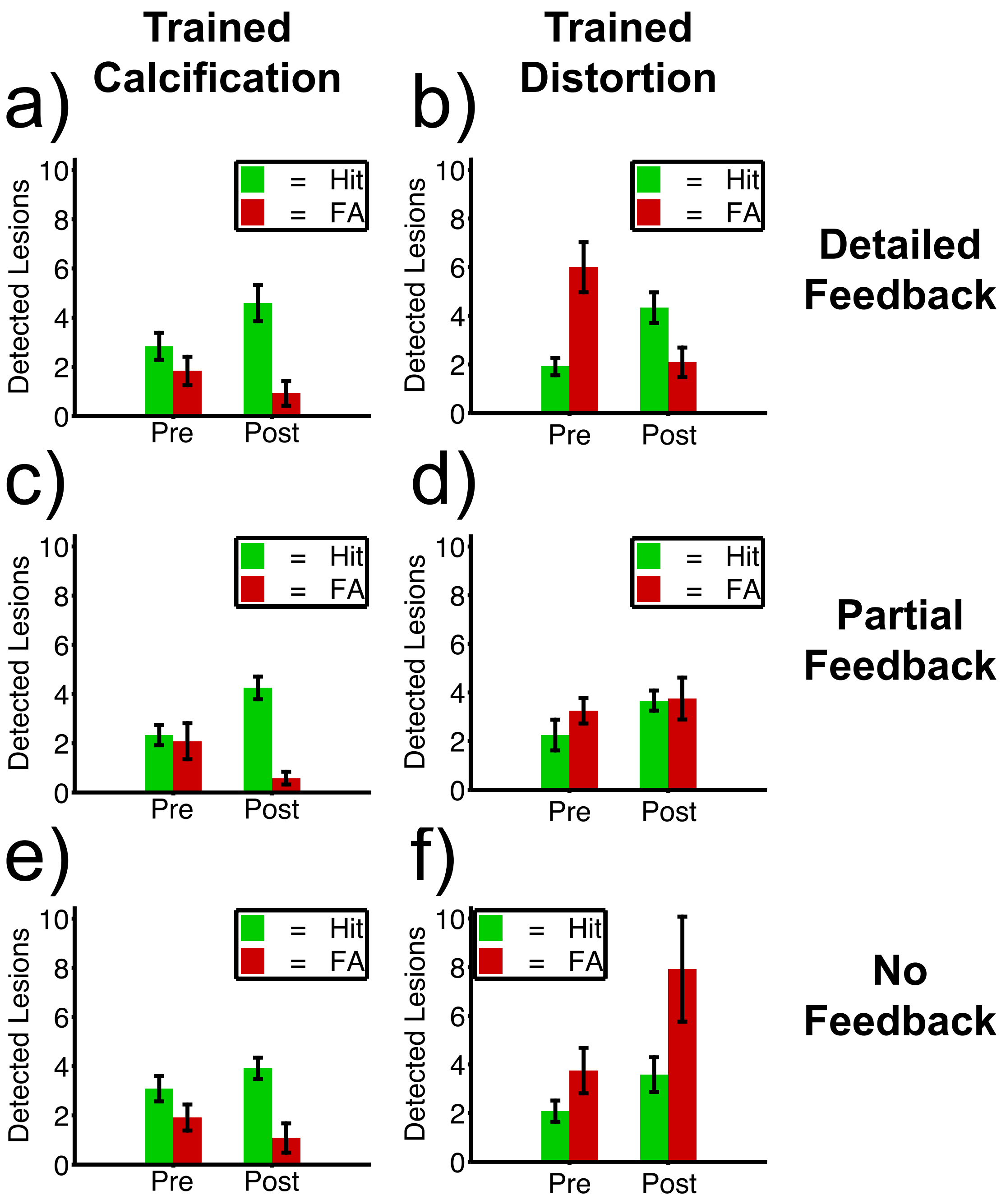


**Supplementary Figure 2**

The same data depicted in Figure 4 but separately for hits (green) and false alarms (red). In each of the test sessions, a total of 10 mammograms with the trained type of lesion were presented alongside 10 mammograms with the untrained type of lesion and 30 normal mammograms. The y-axis shows the mean (± SE) number of correctly (hit) and incorrectly (false alarm) detected trained type of lesion. (a) and (b): with detailed feedback during training. (c) and (d): with partial feedback during training. (e) and (f): with no feedback during training.

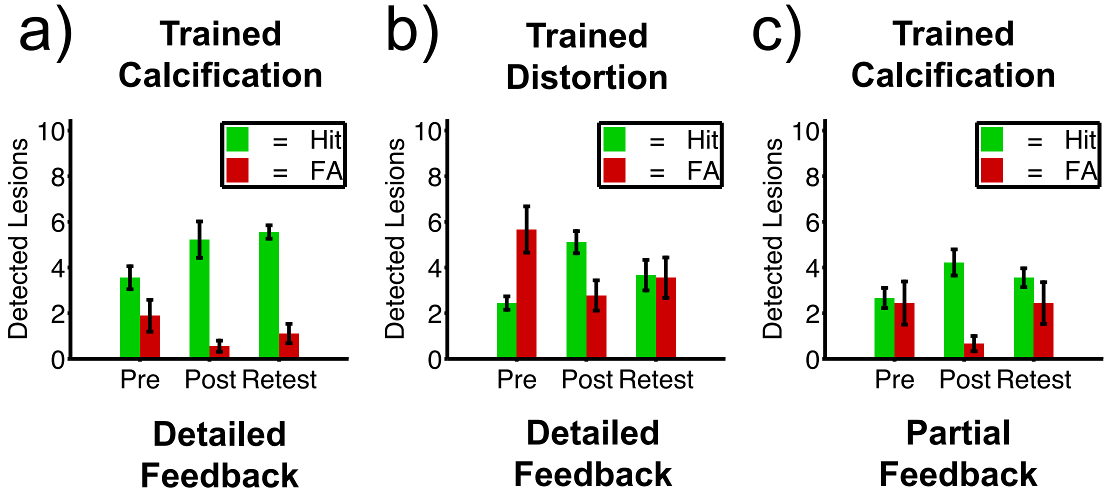


**Supplementary Figure 3**

The same data depicted in Figure 5 but separately for hits (green) and false alarms (red). In each of the test sessions, a total of 10 mammograms with the trained type of lesion were presented alongside 10 mammograms with the untrained type of lesion and 30 normal mammograms. The results are shown for subjects who participated in the retest (n = 9 subjects in the detailed and n = 9 subjects in the partial feedback training conditions). The y-axis shows the mean (± SE) number of correctly (hit) and incorrectly (false alarm) detected trained type of lesion. (a): subjects trained on calcifications with detailed feedback during training. (b): subjects trained on distortions with detailed feedback during training. (c): subjects trained on calcifications with partial feedback during training.

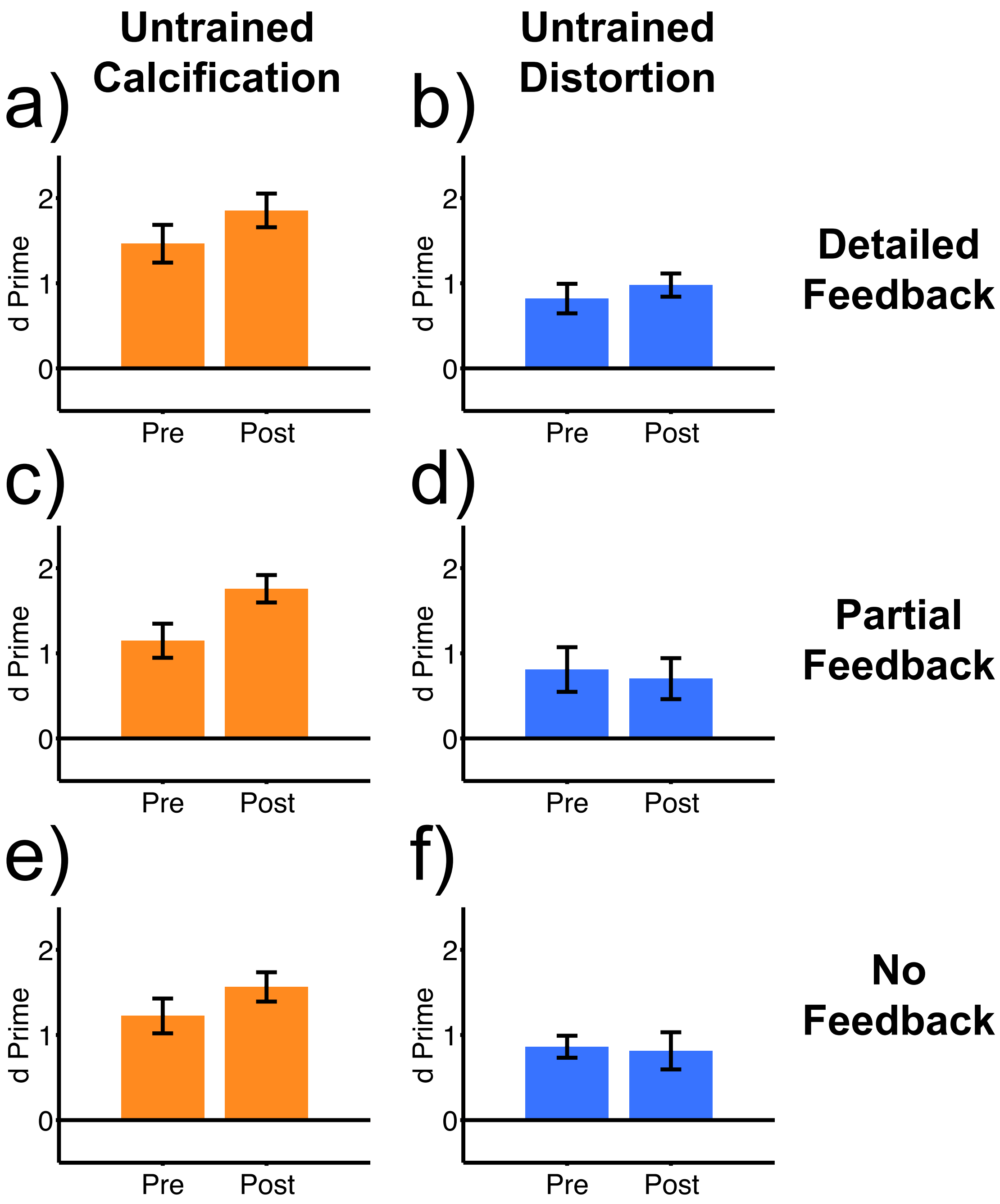


**Supplementary Figure 4**

PL for untrained lesions. Pre and Post on the x-axis correspond to the pre- and posttest, which were conducted before and after training, respectively. Orange bars represent the mean (± SE) observer sensitivity (d') for untrained calcifications in subjects trained on distortions (n = 12 for each feedback condition). Blue bars represent the mean (± SE) observer sensitivity for untrained distortions in subjects trained on calcifications (n = 12 for each feedback condition). (a) and (b): with detailed feedback during training. (c) and (d): with partial feedback during training. (e) and (f): with no feedback during training.

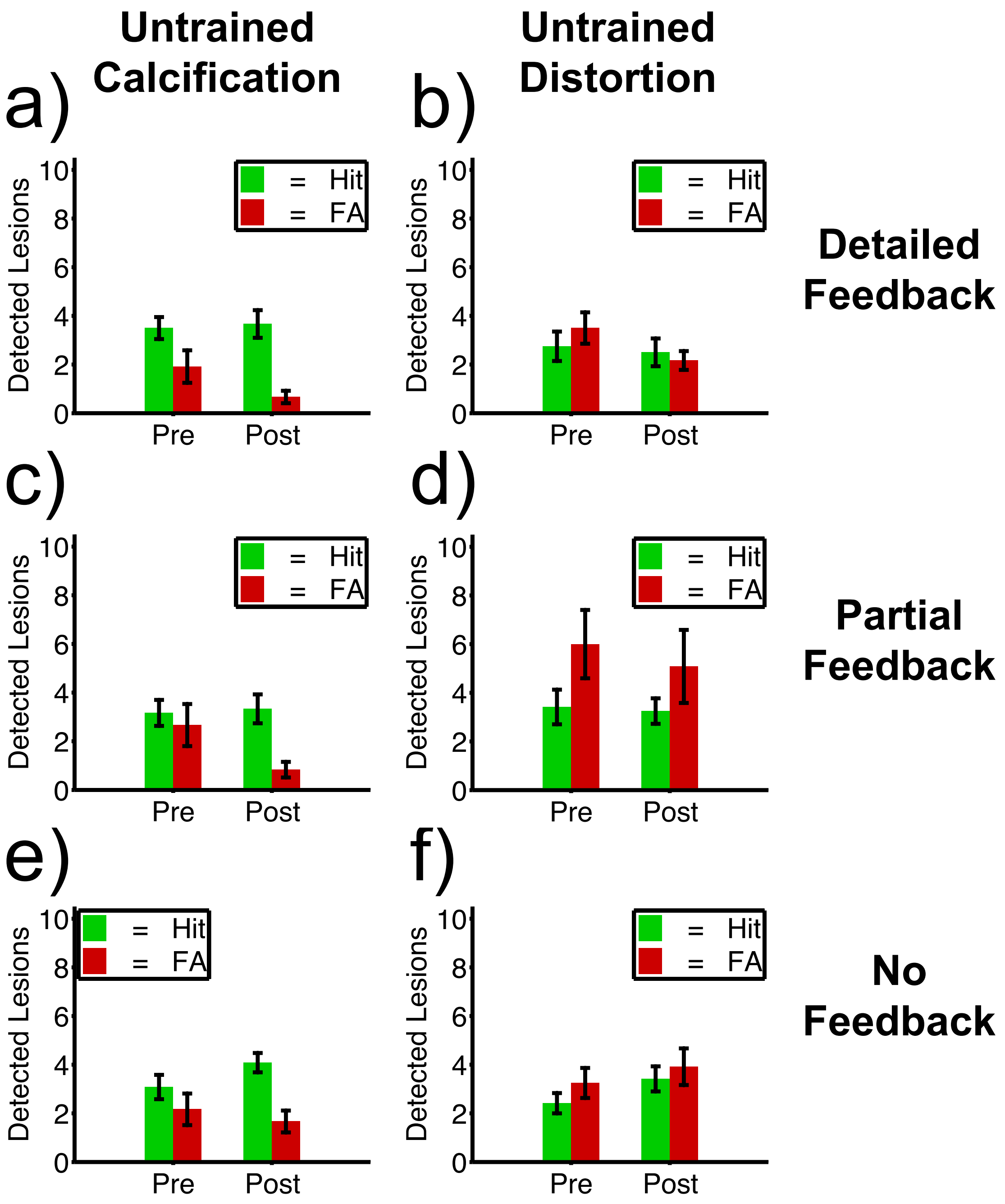


**Supplementary Figure 5**

The same data depicted in Supplementary Figure 4 but separately for hits (green) and false alarms (red). In each of the test sessions, a total of 10 mammograms with the untrained type of lesion were presented alongside 10 mammograms with the trained type of lesion and 30 normal mammograms. The y-axis shows the mean (± SE) number of correctly (hit) and incorrectly (false alarm) detected untrained type of lesion. (a) and (b): with detailed feedback during training. (c) and (d): with partial feedback during training. (e) and (f): with no feedback during training.

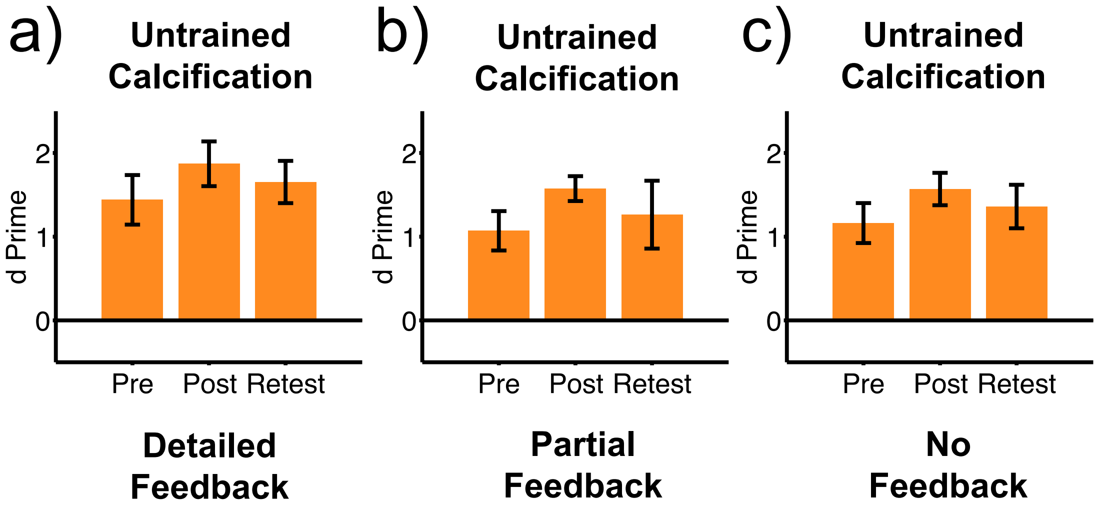


**Supplementary Figure 6**

Retention of PL for untrained calcifications. Pre and Post on the x-axis represent the pretest and posttest, respectively. The retest was conducted six months after the posttest. Orange bars represent the mean (± SE) observer sensitivity (d') for untrained calcifications in subjects who were trained on distortions and participated in the retest (n = 9 subjects for detailed feedback, n = 6 subjects for partial feedback, n = 10 subjects for no feedback). (a): with detailed feedback during training. (b): with partial feedback during training. (c): with no feedback during training.

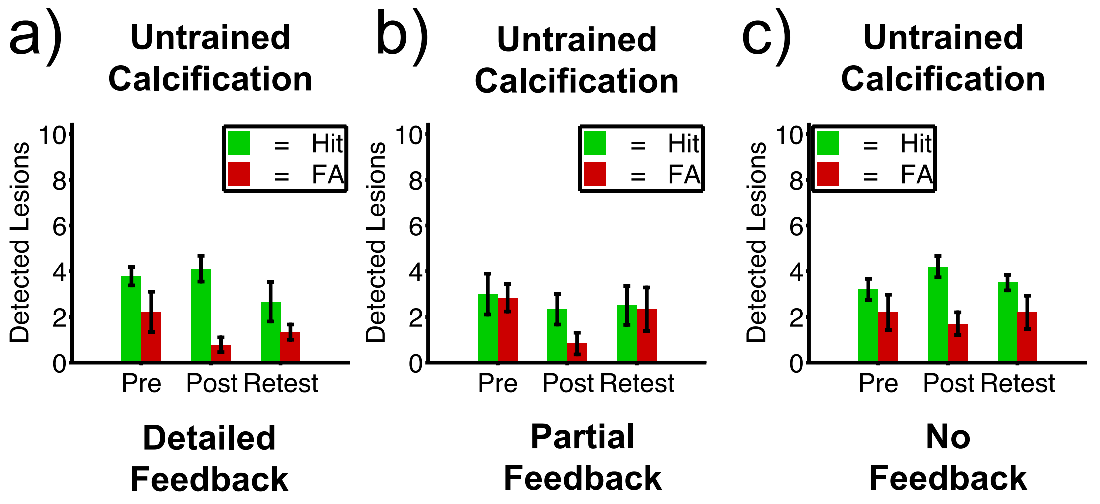


**Supplementary Figure 7**

The same data depicted in Supplementary Figure 6 but separately for hits (green) and false alarms (red). In each of the test sessions a total of 10 mammograms with the untrained type of lesion were presented alongside 10 mammograms with the trained type of lesion and 30 normal mammograms. The results are shown for subjects who trained on distortions and participated in the retest. The y-axis shows the mean (± SE) number of correctly (hit) and incorrectly (false alarm) detected untrained calcifications. (a): with detailed feedback during training. (b): with partial feedback during training. (c): with no feedback during training.
